# Resistance of endothelial cells to SARS-CoV-2 infection *in vitro*

**DOI:** 10.1101/2020.11.08.372581

**Authors:** Blerina Ahmetaj-Shala, Thomas P. Peacock, Laury Baillon, Olivia C. Swann, Hime Gashaw, Wendy S. Barclay, Jane A. Mitchell

## Abstract

**Rationale:** The secondary thrombotic/vascular clinical syndrome of COVID-19 suggests that SARS-CoV-2 infects not only respiratory epithelium but also the endothelium activating thrombotic pathways, disrupting barrier function and allowing access of the virus to other organs of the body. However, a direct test of susceptibility to SARS-CoV-2 of authentic endothelial cell lines has not been performed.

**Objective:** To determine infectibility of primary endothelial cell lines with live SARS-CoV-2 and pseudoviruses expressing SARS-CoV-2 spike protein.

**Methods and Results:** Expression of ACE2 and BSG pathways genes was determined in three types of endothelial cells; blood outgrowth, lung microvascular and aortic endothelial cells. For comparison nasal epithelial cells, Vero E6 cells (primate kidney fibroblast cell line) and HEK 293T cells (human embryonic kidney cells) transfected with either ACE2 or BSG were used as controls. Endothelial and Vero E6 cells were treated with live SARS-CoV-2 virus for 1 hour and imaged at 24 and 72 hours post infection. Pseudoviruses containing SARS-CoV-2, Ebola and Vesicular Stomatis Virus glycoproteins were generated and added to endothelial cells and HEK 239Ts for 2 hours and infection measured using luminescence at 48 hours post infection. Compared to nasal epithelial cells, endothelial cells expressed low or undetectable levels of ACE2 and TMPRSS2 but comparable levels of BSG, PPIA and PPIB. Endothelial cells showed no susceptibility to live SARS-CoV-2 or SARS-CoV-2 pseudovirus (but showed susceptibility to Ebola and Vesicular Stomatitis Virus). Overexpression of ACE2 but not BSG in HEK 239T cells conferred SARS-CoV-2 pseudovirus entry. Endothelial cells primed with IL-1ß remained resistant to SARS-CoV-2.

**Conclusion:** Endothelial cells are resistant to infection with SARS-CoV-2 virus, in line with relatively low levels of ACE2 and TMPRSS2, suggesting that the vascular dysfunction and thrombosis seen in severe COVID-19 is a result of factors released by adjacent infected cells (e.g. epithelial cells) and/or circulating, systemic inflammatory mediators.

## Introduction

COVID-19 represents one of the most important clinical challenges the scientific community has faced in recent memory. In the absence of an effective vaccine and no clear evidence that prior infection confers long-term immunity^1^, there is an urgent unmet need to understand disease pathology and to establish therapeutics that mitigate COVID-19 severity and reduce associated lethality^2^. SARS-CoV-2 spike protein binds to host cells via ACE2^3^ and viral entry is facilitated by the cell-surface protease TMPRSS2^3^ or lysosomal cysteine proteases cathepsin B/L (CTSB, CTSL)^3^. It has also been suggested that, as an alternative pathway, SARS-CoV-2 binds to cells via BSG (Basigin; also known as CD147 or EMMPRIN)^4, 5^, although firm evidence for BSG as a standalone receptor for SARS-CoV-2 remains the subject of investigation with a recent study noting no ‘direct’ binding of SARS-CoV-2 spike protein to BSG^6^.

Initial infection with SARS-CoV-2 occurs via the respiratory epithelium^7^. In most people symptoms are mild but in a significant minority COVID-19 progresses to severe disease and in those that ‘recover’ symptoms can persist leading to a syndrome recently defined as ‘long-COVID’^8^. In severe COVID-19 and in long-COVID multiple organs including the cardiovascular system are affected^9, 10^. This secondary thrombotic/vascular clinical syndrome of COVID-19 suggests that SARS-CoV-2 infects not only respiratory epithelium but also the endothelium disrupting barrier function and allowing dissemination to other organs of the body^11^. This notion is supported by early reports showing that SARS-CoV-2 infects endothelial cells *in vitro*^12^ and *in vivo^13^*. However, for the *in vitro* studies performed to date, non-physiological cell lines were used and for *in vivo* studies serious concerns have been raised regarding the validity of the interpretation of data and the resulting conclusions^14^. Currently, therefore, it is not clear if endothelial cells are permissive to SARS-CoV-2 or not. Only by addressing this can biomarkers, mechanisms and therapeutic targets directed at the vasculopathy and thrombosis associated with COVID-19 be established.

We have recently shown that blood outgrowth endothelial cells express relatively low levels of ACE2 and TMPRSS2 but high levels of BSG and speculated that if endothelial cells are infected by SARS-CoV-2, BSG, rather than ACE2, would act as the receptor for viral entry^15^. However, in our previous work and that of others, a direct test of susceptibility to SARS-CoV-2 of authentic endothelial cell lines has not been performed.

Here we have performed the first infection study using primary endothelial cell lines with live SARS-CoV-2 and pseudoviruses (PV) expressing SARS-CoV-2 spike protein. Specifically, we used (i) blood outgrowth endothelial cells, selected for their utility as biomarkers and in personalised medicine, (ii) lung microvascular endothelial cells, selected based on likely points of access of the virus and (ii) aortic endothelial cells, selected as representative of systemic vessels. For comparison we used positive control cells including nasal epithelial cells, Vero E6 cells (primate kidney fibroblast cell line) and HEK 293T cells (human embryonic kidney cells) transfected with either ACE2 or BSG.

## Methods

### Biosafety statement

All work performed was approved by the local genetic manipulation (GM) safety committee of Imperial College London, St. Mary’s Campus (centre number GM77), and the Health and Safety Executive of the United Kingdom, under reference CBA1.77.20.1.

### Cells

Primary human endothelial cell lines including blood outgrowth endothelial cells (obtained in house^16^), lung microvascular endothelial cells and aortic endothelial cells (obtained from Lonza; UK) were used in this study. In each case cells from 3 separate donors/ per endothelial cell type were used except for qPCR studies using blood outgrowth endothelial cells where cells from 2 donors were used. Nasal epithelial cells were obtained from Promocell (Germany) and grown in submersed culture and differentiated nasal epithelial cells (MucilAir™) were obtained from Epithelix (Switzerland) and grown in air-liquid interface culture.

African green monkey (Vero E6) cells (ATCC) and human embryonic kidney cells (293T) - 293T(ATCC) were cultured in Dulbecco’s modified Eagle’s medium (DMEM), supplemented with 10% foetal bovine serum (FBS), 1% non-essential amino acids (NEAA) and 1% penicillin-streptomycin (P/S; Gibco). HEK 293T-ACE2 cells were produced as previous described^17^ (Figure 3, Supplementary Figure 2) and maintained in HEK 293T media supplemented with 1 μg/ml of puromycin. For transient transfections HEK 293T cells seeded in 10cm^2^ plates were transfected with 1 μg of empty pCAGGs vector, pCAGGs-ACE2-FLAG, or pCAGGS-BSG (synthesised by GeneArt; ThermoFisher), 24 hours later cells were transferred to 96 well plates or lysed for western blot analysis. Aortic and blood outgrowth endothelial cells were maintained on uncoated plates in Endothelial Cell Growth Medium-2 (EGM-2; Lonza, UK), while lung microvascular cells were maintained on uncoated plates in Microvascular Endothelial Cell Growth Medium-2 (EGM-2 MV; Lonza, UK). Nasal epithelial cells were maintained in Airway Epithelial Growth Media (Promocell, Germany) while MucilAir™ nasal epithelial cells were maintained in MucilAir Culture Medium (Epithelix, Switzerland). All media for endothelial and epithelial cells were supplemented with their appropriate BulletKits, 10% FBS (Biosera, UK) and 1% P/S (Gibco, UK). Treatment protocols (except for those using MucilAir™ cells) were conducted using ‘treatment media’ (EGM-2 medium (Promocell, Germany) supplemented with 2% FBS (Promocell, Germany) and 1% P/S (Gibco, UK). For all experiments, endothelial and nasal epithelial cells were used between passage 4-6. All cells were maintained at 37°C, 5% CO_2_.

### RT-qPCR

Endothelial and nasal epithelial cells were plated in duplicate wells on uncoated 6 well plates in their cell-specific media (see above) and grown to confluence. The day before RNA extraction media was replaced with the treatment media. After 24 hours cells were washed with phosphate-buffered saline (PBS), duplicate wells combined and RNA extracted using RNeasy Extraction kit (Qiagen, UK). RNA was converted to cDNA using the iScript cDNA synthesis kit (BioRad, CA, USA). Gene expression levels were determined using a TaqMan expression assay, with the following primers (ThermoScientific, UK); ACE2 (Hs01085333_m1), TMPRSS2 (Hs00237175_m1), BSG (Hs00936295_m1), PPIA (Hs04194521_s1) and PPIB (Hs00168719_m1). Genes were quantified relative to housekeeping genes (GAPDH and 18S) by the comparative Ct method.

### SARS-CoV-2 live virus infection studies

For infection studies with Mucilair human airway epithelial cells, SARS-CoV-2/England/IC19/202 (IC19)^18^ was diluted in serum-free DMEM, 1% NEAA, 1% P/S to a multiplicity of infection (MOI) of 0.1. Inoculum was added to the exposed apical face of the cells and incubated at 37°C for 1 hour. Inoculum was then removed and cells maintained as described above. At each timepoint in the infection, virus was collected from the apical surface of the cultures by washing with pre-warmed PBS for 10 minutes at 37°C and quantified by plaque assay on Vero E6 cells. Briefly, cells were washed with PBS then serial dilutions of inoculum, diluted in serum-free DMEM, 1% NEAA, 1% P/S, were overlayed onto cells for one hour at 37°C. Inoculum was then removed and replaced with SARS-CoV-2 overlay media (1x minimal essential media (MEM), 0.2% w/v bovine serum albumin, 0.16% w/v NaHCO3, 10mM Hepes, 2mM L-Glutamine, 1x P/S, 0.6% w/v agarose). Plates were incubated for 3 days at 37°C before overlay was removed and cells were stained for 1 hour at RT in crystal violet solution.

For infection studies using endothelial cells, cells were plated on sterile round 16mm diameter coverslips in 12 well plates (5×10^4^ cells/well) without coating. Their usual media (see above) was added and cells allowed to settle overnight. Control Vero E6 cells were plated to achieve approximately 80% confluency. The following day, media was removed and cells washed in PBS. For endothelial cells, treatment media was added either alone (untreated/ control) or with IL-1β (10ng/ml) for 3 hours. Meanwhile IC19^18^ was diluted in serum free DMEM, 1%NEAA, 1% P/S, to a multiplicity of infection (MOI) of 0.1. After 3 hours treatment with either media alone or IL-1β, media was replaced with IC19 containing inoculum and incubated at 37°C for 1 hour. Inoculum was then removed and replaced with treatment media and cells were maintained until 24, 48 or 72 hours post-infection. At the appropriate timepoint, treatment media was removed and cells were fixed in 4% paraformaldehyde (PFA) for 30 minutes. PFA was removed with three washes of PBS. Cover slips were dehydrated in an ethanol series and stored in 100% ethanol at −20°C until further processing.

### Fluorescent imaging

For fluorescent imaging, infected cells on cover slips were first rehydrated in an ethanol series. Cells were permeabilised in PBS with 0.5% triton-X for 10 minutes, washed 3x in PBS, then blocked in PBS with 2% bovine serum albumin (BSA) and 0.1% tween (blocking buffer). Primary antibodies were diluted 1:1000 in blocking buffer and incubated on cells at room temperature for 1 hour. Primary antibodies used were against spike protein (S) (Mouse monoclonal, Gene tex (1A9)) or nucleoprotein (N) (Rabbit monoclonal, Sino Biological). Following 3x PBS washes, cells were incubated with secondary antibodies (anti-rabbit 488, anti-mouse 594) diluted 1:500 and DAPI (diluted 1 in 1000) in PBS with 2% BSA at room temperature for 1 hour in the dark. Stained cover slips were mounted on glass slides in ProLong gold antifade mounting medium (Invitrogen). Images were acquired using either a Zeiss Axiovert 135 TV microscope (Figure 2B) or a Zeiss Cell Observer widefield microscope (Figure 2C). All images were analysed and prepared using FIJI software^19^. For each cell type/condition slides were first reviewed by eye before representative fields of view were captured. Images were captured and processed in an identical manner across each experiment to ensure fair comparison either as a single plane of focus (Figure 2B) or as Z stacks presented as maximum intensity projections (Figure 2C). For quantitative analysis, all images were blinded and independently scored between 1-5, where 1= 0-2, 2= 3-5, 3=6-8, 4= 9-10 and 5= >10 nucleocapsid stained cells.

### Lentiviral pseudovirus studies

Lentiviral pseudotypes were generated as previous described^17^. Briefly, dishes of HEK 293T cells were co-transfected with 1 μg of HIV packaging plasmid, pCAGGs-GAGPOL, 1.5 μg of luciferase reporter genome construct, pCSLW, and 1 μg of the named envelope proteins in pcDNA3.1. Media was replaced at 24 hours to remove transfection reagents and plasmids and then pseudovirus-containing supernatants were harvested at 48 and 72 hours post-transfection. Pooled supernatant was then passed through a 0.45 μm filter, aliquoted and frozen at −80°C.

Cells were plated in duplicate wells in 96 well plates (1×10^4^/ well) in their own media and allowed to settle overnight in their appropriate growth media. The next day media was removed and replaced with treatment media alone (untreated/ control) or with IL-1β (10ng/ml) for 3 hours. Media was then removed and cells were treated with pseudovirus containing-supernatants added for 2 hours before replacing with treatment media. Cells were then left for 48 hours before lysis with cell culture lysis buffer (Promega). Luminescence was read on a FLUOstar Omega plate reader (BMF Labtech) using the Luciferase Assay System (Promega).

### Western Blot

HEK 293Ts transfected with different potential receptors or empty vector were lysed 24 hours post-transfection with RIPA buffer (150mM NaCl, 1% NP-40, 0.5% sodium deoxycholate, 0.1% SDS, 50mM TRIS, pH 7.4) supplemented with an EDTA-free protease inhibitor cocktail tablet (Roche). Cell lysates were then mixed with 4x Laemmli sample buffer (Bio-Rad) with 10% β-mercaptoethanol. Protein samples were run on a 4-15% mini-PROTEAN TGX SDS-PAGE gel (Bio-Rad) and transferred onto a nitrocellulose membrane by semi-dry transfer. Membranes were probed with the primary antibodies: mouse anti-FLAG (Sigma: F1804); mouse anti-tubulin (abcam; ab7291); and rabbit anti-CD147 (abcam; ab108308). Near infra-red (NIR) secondary antibodies, IRDye® 680RD Goat anti-mouse (abcam; ab216776) and IRDye® 800CW Goat anti-rabbit (abcam; ab216773)) were subsequently used to detect primary antibodies. Western blots were visualised using an Odyssey Imaging System (LI-COR Biosciences).

### IL-6 and IL-8 ELISA

IL-6 and IL-8 was measured using duo set ELISAs from R&D Systems according to manufactures instructions.

### Data analysis

All data were analysed on GraphPad Prism v8 and are shown as individual data points and/or mean +/− standard error of the mean (SEM) for samples as described in the figure legends.

## Results

In our recent study, where we performed a systematic analysis of online transcriptomic databases in endothelial cells and nasal and bronchial epithelium, we found endothelial cells express relatively low levels of ACE2 and TMPRSS2 but high levels of BSG and PPIA/PPIB (also known as cyclophilin A/B; CypA/B)^15^. PPIA and PPIB are potential co-ligands for pathogen entry to cells via the BSG pathway. Here we have validated and extended our earlier studies by measuring gene expression using qPCR in blood outgrowth endothelial cells, lung microvascular endothelial cells, aortic endothelial cells and, for comparisons, nasal epithelium. In line with our previous observations, compared to nasal epithelial cells, endothelial cells expressed low or undetectable levels of ACE2 and TMPRSS2 but comparable levels of BSG, PPIA and PPIB, (Figure 1; Supplementary Table 1). These observations suggest that if endothelial cells are suspectable to infection with SARS-CoV-2 it may be occur in an ACE2-independent pathway, for example via the putative BSG receptor pathway.

**Figure 1:**
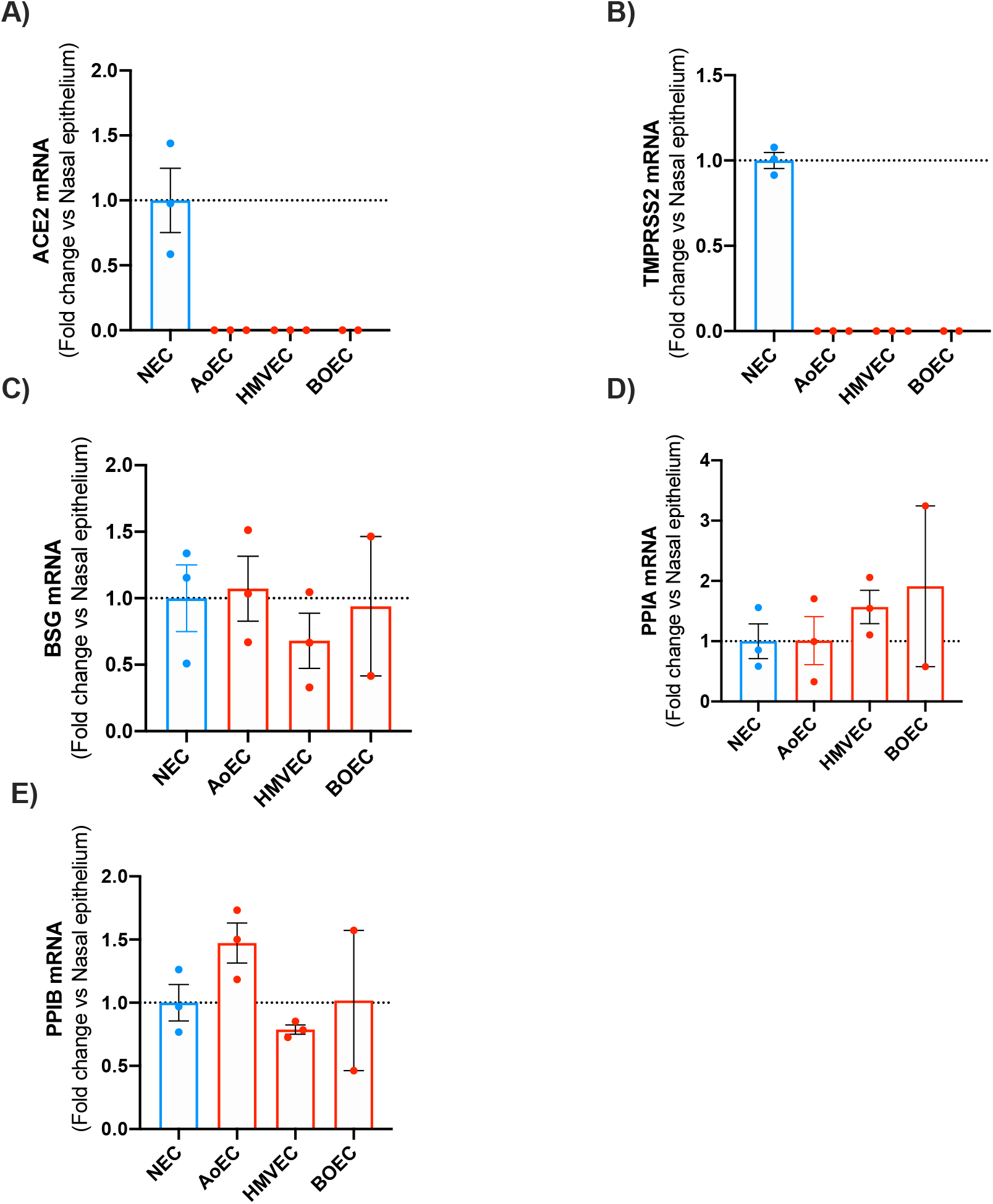
mRNA expression of *ACE2*, *TMPRSS2*, *BSG*, *PPIA* and *PPIB* in human nasal epithelial cells (NEC) and endothelial cells (aortic, microvascular and blood outgrowth). Expression levels for the genes *ACE2*, *TMPRSS2*, *BSG*, *PPIA* and *PPIB* were obtained from aortic (AoEC), microvascular (HMVEC) and blood outgrowth (BOEC) endothelial cells and nasal epithelial cells (NEC). Data for each donor were normalised using the average of the housekeepers (18S and Gapdh) and analysed using a comparative Ct method (2ΔΔCt). Data are shown as the mean +/− SEM fold change compared to nasal epithelium (n=3 wells using cells from 2 donors) for AoEC (n=3 wells using cells of 3 separate donors), HMVEC ((n=3 wells using cells of 3 separate donors and BOECs (n=2 wells using cells of 2 separate donors).

Next, using experimental conditions where nasal epithelial cells grown in liquid: air interface culture were highly susceptible to infection with SARS-CoV-2 (Figure 2A), we inoculated endothelial cells and, in parallel, Vero E6 cells, with live SARS-CoV-2. Initially we performed pilot studies using blood outgrowth endothelial cells and Vero E6 cells incubated for 24 hours after a 1 hour inoculation (Figure 2B). These were followed by separate experiments using each of the 3 endothelial cell types incubated for up to 72 hours post inoculation with live SARS-CoV-2 (Figure 2C). In both studies by indirect immunofluorescence microscopy (IF) we could see cells staining positive for viral antigens (nucleocapsid protein or spike) in Vero E6 cells, indicating virus replication, but not in the endothelial cell types at any of the time points tested (Figures 2B and C; Supplementary Figure 1). These results suggest that endothelial cells have a block to SARS-CoV-2 infection, however it is not clear whether this block is at virus entry or upon a post-entry step in replication.

**Figure 2:**
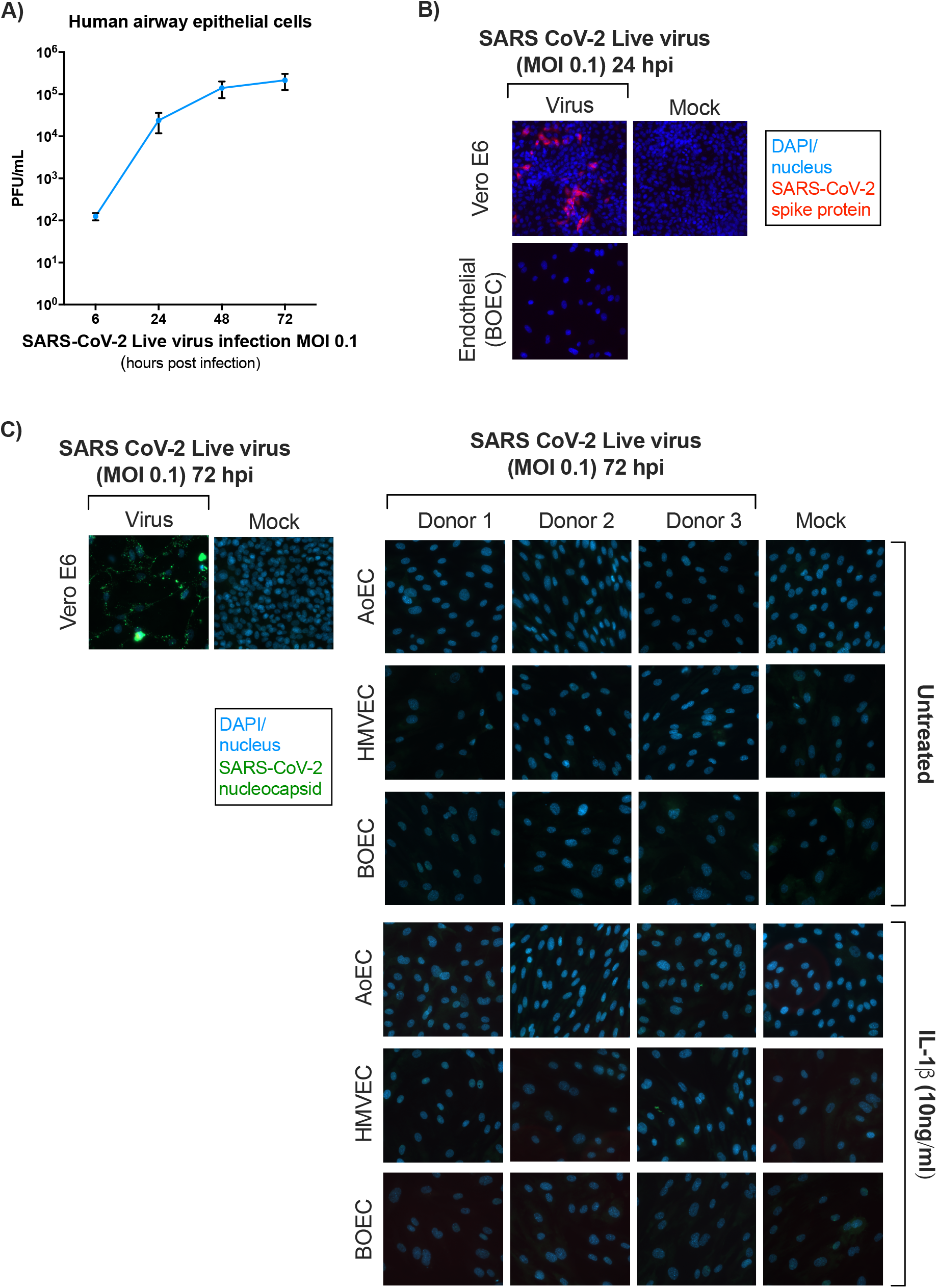
SARS-CoV-2 live virus infection in human airway epithelial cells in air:liquid interface, Vero E6 and endothelial cells. Human airway epithelial cells grown in an air:liquid interface environment from MucilAir™ were infected with SARS-CoV-2 live virus (MOI 0.1) and the plaque forming units (measure of infectious virus obtained from permissive cells) was determined over time (6, 24, 48 and 72 hours post-infection). In separate studies, the levels of SARS-CoV-2 nucleocapsid or spike protein in Vero E6 and endothelial cells (treated with media only (untreated) or IL-1ß (10ng/ml; 3hrs)) at 24 (B) and 72 (C) hours post infection with SARS-CoV-2 (MOI 0.1) were determined using florescent imaging. Mock controls (media only) experiments were run simultaneously using each endothelial cell line. Data is shown as n=3 (pooled donors) for Mucilair cells (A) and n=3 (separate donors) for human aortic (AoEC), lung microvascular (HMVEC) and blood outgrowth endothelial cells (BOEC). Data are shown are mean +/− SEM for panel A and representative images shown for panel A and representative images for panels B and C.

To investigate whether the block to virus infection was at entry we performed studies using pseudoviruses based on lentiviral vectors, expressing SARS-CoV-2 spike protein and, by way of positive controls for viral infectability, pseudovirus expressing the glycoproteins of Vesicular Stomatitis Virus (VSV-G), which shows broad cell-type tropism and Ebola, which has previously been shown to efficiently infect endothelial cells^20^. As with the virus infection, endothelial cells were not susceptible to SARS-CoV-2 spike-mediated entry, showing similar levels of luciferase signal as bald pseudovirus expressing no viral glycoprotein (Figure 3A), but were found to be susceptible to pseudovirus expressing Ebola glycoprotein and VSV-G (Figure 3B and D). Furthermore, HEK 293T cells stably expressing ACE2 showed high susceptibility to all 3 pseudoviruses. These observations are in line with low level ACE2 expression across our endothelial cell lines and suggest that the high levels of BSG expression do not compensate to allow infection with SARS-CoV-2. To address this assumption, we investigated SARS-CoV-2 pseudovirus entry into HEK 293T cells transiently transfected with either ACE2 or BSG (successful transfection was confirmed using western blot analysis; Supplementary Figure 2). From this data it was clear that transfected ACE2 but not BSG efficiently mediated SARS-CoV-2 pseudovirus entry (Figure 3D).

**Figure 3:**
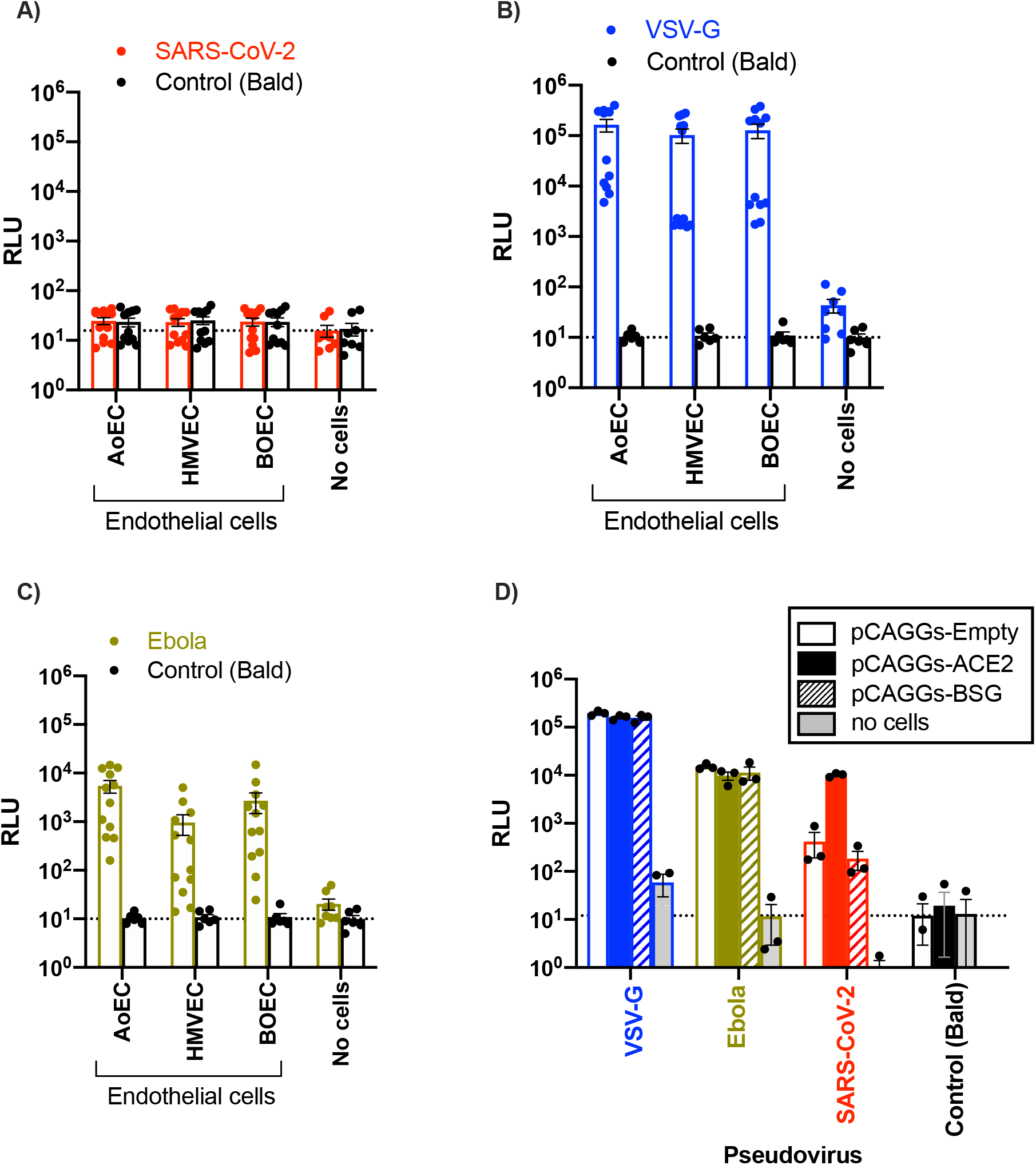
Pseudovirus infection with SARS-CoV-2 (A), vesicular stomatitis virus G (VSV-G) (B), Ebola (C) in endothelial cells or SARS-CoV-2 in kidney embryonic kidney (HEK 293T) cells transfected with ACE2 or BSG (D). SARS-CoV-2 (A) and VSV-G and Ebola (B) pseudovirus levels were measured using a Luciferase Assay System 48 hours post-transduction. Data were from n=12 wells using cells from 3 donors separate donors in experiments performed on 2 separate occasions for aortic (AoEC), microvascular (HMVEC) and blood outgrowth (BOEC) (A-C); ACE2-HEK-2932T and BSG-HEK-2932T are n=3. Data are expressed as individual values and as mean +/− SEM. Dotted lines on graphs indicate background signal as determined by the mean of control/bald (no cells or empty).

Finally, to investigate how an inflammatory environment might facilitate virial entry of SARS-CoV-2 into endothelial cells additional protocols were performed where cells were primed with IL-1ß. At 24 hours IL-1ß primed endothelial cells released increased levels of IL-6 and IL-8, demonstrating a positive inflammatory response (Supplementary Figure 3). However, under these conditions each of the endothelial cell types remained resistant to SARS-CoV-2 spike mediated entry (Figure 4A and B).

**Figure 4:**
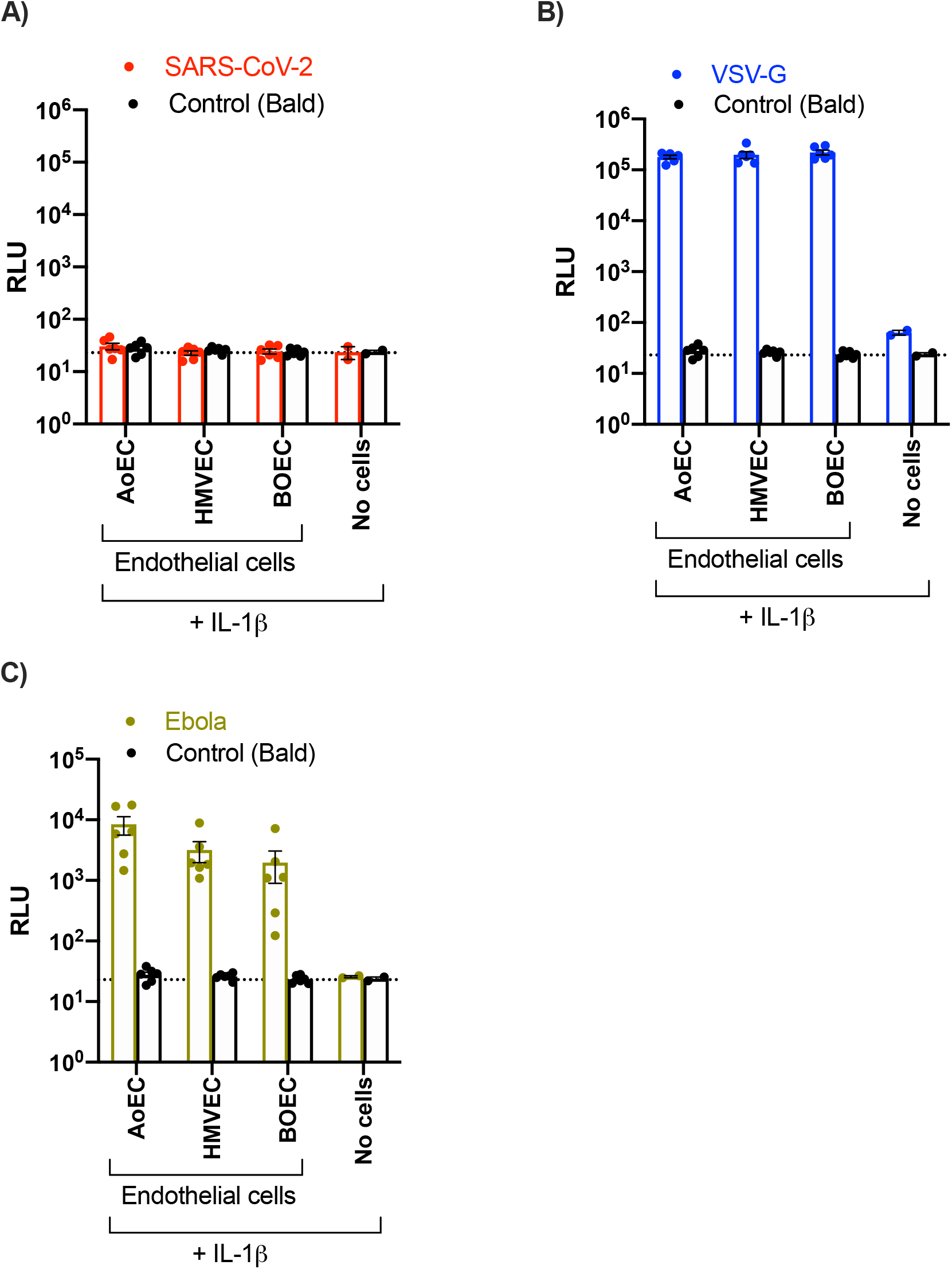
Susceptibility of endothelial cells treated with IL-1β to pseudoviruses expressing SARS-CoV-2 spike (A), vesicular stomatitis virus glycoprotein (VSV-G) (B) or Ebola glycoprotein (C). SARS-CoV-2 (A) and VSV-G and Ebola (B) pseudovirus entry was measured using a Luciferase Assay System 48 hours post infection in endothelial cells treated with IL-1β (10 ng/ml) for 3 hours prior to infection. Data were from n=6 wells from n=3 donors of endothelial cells; aortic (AoEC), microvascular (HMVEC) and blood outgrowth (BOEC). Data are expressed as individual values and mean +/− SEM. Dotted lines on graphs indicate background signal as determined by the mean of control/bald (no cells).

## Discussion

There is mounting speculation that the vascular and thrombotic sequela associated with severe COVID-19 is a result of endothelial cell infection with SARS-CoV-2. However, there is currently no direct evidence to support this idea. In response to this, we have used standard virology protocols to directly address the question of *‘are endothelial cells permissive to infection with SARS-CoV-2 virus?*’ We have used 3 carefully selected primary endothelial cells lines from 3 separate donors for each line. The results were unambiguous and interpreted alongside data from robust positive control cells infect with live virus and pseudoviruses. Taken together our data suggest that endothelial cells are not susceptible to infection with SARS-CoV-2 because they express insufficient levels of ACE2. Our findings that endothelial cells remain non-susceptible to infection with SARS-CoV-2 despite expressing high levels of BSG suggests that the BSG pathway is not a functional entry pathway for SARS-CoV-2 in endothelial cells. This idea is confirmed from our studies using HEK 293T cells transiently transfected with BSG which, in contrast to ACE2, did not make cells susceptible to SARS-CoV-2 spike-mediated entry.

Our results showing that endothelial cells are not susceptible to the SARS-CoV-2 virus suggests that the vascular dysfunction and thrombosis seen in severe COVID-19 is a result of factors released by adjacent infected cells (e.g. epithelial cells) and/or circulating, systemic inflammatory mediators. In addition, it is possible that SARS-CoV-2 virus may act as an inflammatory stimulus either via the spike protein or other structural components without the requirement for conventional binding, cell entry or virus replication^21^. Our study does not rule out this possibility and this remains the subject of further investigation. Our work also suggests that where viremia occurs in COVID-19, SARS-CoV-2 passes through the endothelium facilitated by loss of barrier function, as a result of local inflammation at the site of infection.

While we have worked to the highest standards with empirical virology protocols, there are limitations in our approach, and we cannot definitively conclude that endothelial cells are non-susceptible to infection by SARS-CoV-2 in some individuals or in some highly specific conditions *in vivo.* This is because the assay systems we used did not take account of critical factors present in at risk populations and/or at the site of inflammation. In an attempt to take account of basic inflammatory conditions we performed experiments in cells primed with IL-1ß, which did not confer infectability to any of our endothelial lines. However, as we find more about the complex mix of inflammatory mediators present in the lung and circulation in COVID-19 and the specific biological factors that predispose certain groups of individuals to severe disease, these can be recapitulated within *in vitro* assay systems. Nonetheless what our study does prove is that if endothelial cells are susceptible to SARS-CoV-2 at some distant point in the natural history of COVID-19, the pathways of viral entry are more complex than for airway epithelium.

## Acknowledgements

We thank the FILM Facility at Imperial College London for the use of the microscopes and their technical support.

## Supplementary Tables and Figures

**Supplementary Table 1:** Average raw Ct values from qRT-PCR. Expression levels (Ct) for the genes *ACE2*, *TMPRSS2*, *BSG*, *PPIA* and *PPIB* were obtained from aortic (AoEC), microvascular (HMVEC) and blood outgrowth (BOEC) endothelial cells and nasal epithelial cells (NEC). Data for each donor were corrected using the average of the housekeepers (*18S* and *GAPDH*) and analysed using a comparative Ct method (2ΔΔCt). Data are shown as the mean +/− from n=3 wells using cells from 3 separate donors for AoEC and HMVEC and n=3 wells using cells from 2 separate donors for NEC and n=2 wells from 2 separate for BOECs.

| Gene | Cell type | Raw CT values |  |  |
| --- | --- | --- | --- | --- |
| Donor/well |  | 1 | 2 | 3 |
| *ACE2* | AoEC | 38.3 | 37.5 | 36.2 |
|  | HMVEC | Undet. | Undet. | Undet. |
|  | BOEC | Undet. | 39.3 | N/A |
|  | NEC | 31.3 | 31.2 | 30.7 |
| *TMPRSS2* | AoEC | Undet. | Undet. | Undet. |
|  | HMVEC | Undet. | Undet. | Undet. |
|  | BOEC | Undet. | Undet. | Undet. |
|  | NEC | 29.2 | 27.6 | 27.5 |
| *BSG* | AoEC | 20.2 | 20.3 | 21.2 |
|  | HMVEC | 23.2 | 24.3 | 21.0 |
|  | BOEC | 23.4 | 20.4 | N/A |
|  | NEC | 20.4 | 21.4 | 20.2 |
| *PPIA* | AoEC | 23.7 | 22.0 | 22.5 |
|  | HMVEC | 23.9 | 23.6 | 22.8 |
|  | BOEC | 24.1 | 22.0 | N/A |
|  | NEC | 22.1 | 23.1 | 22.5 |
| *PPIB* | AoEC | 20.6 | 21.0 | 21.2 |
|  | HMVEC | 24.1 | 24.0 | 22.2 |
|  | BOEC | 24.2 | 21.2 | N/A |
|  | NEC | 21.4 | 21.7 | 21.3 |
| *18S/GAPDH* | AoEC | 17.8 | 17.3 | 17.1 |
|  | HMVEC | 19.1 | 20.0 | 17.9 |
|  | BOEC | 21.0 | 16.7 | N/A |
|  | NEC | 18.1 | 17.2 | 17.3 |

**Supplementary Figure 1.**
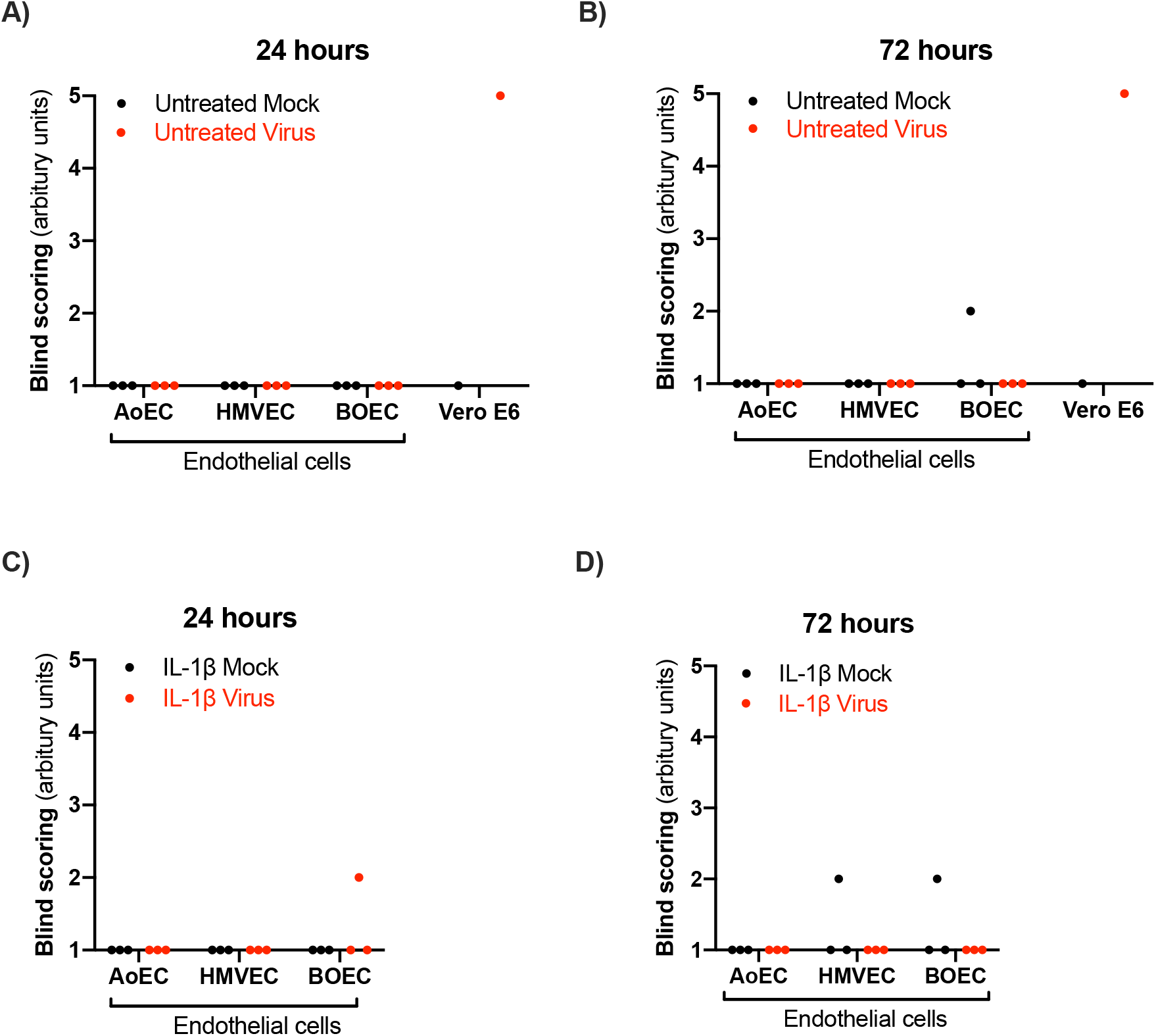
Blind scoring of SARS-CoV-2 live virus infection in Vero E6 and endothelial cells. Levels of SARS-CoV-2 nucleocapsid and spike protein in Vero E6 and endothelial cells at 24 and 72 hours post infection with SARS-CoV-2 (MOI 0.1) in untreated (A-B) and IL-1ß (10ng/ml; 3 hours) (C-D) were determined using florescent imaging. One representative image was taken and blinded images were scored between 1-5, where 1= 0-2, 2= 3-5, 3= 6-8, 4= 9-10 and 5= >10 virus nucleocapsin/spike protein staining. Data are shown as individual scores for n=3 (separate donors) for human aortic (AoEC), lung microvascular (HMVEC) and blood outgrowth endothelial cells (BOEC) and n=1 for Vero E6 cells (untreated).

**Supplementary Figure 2:**
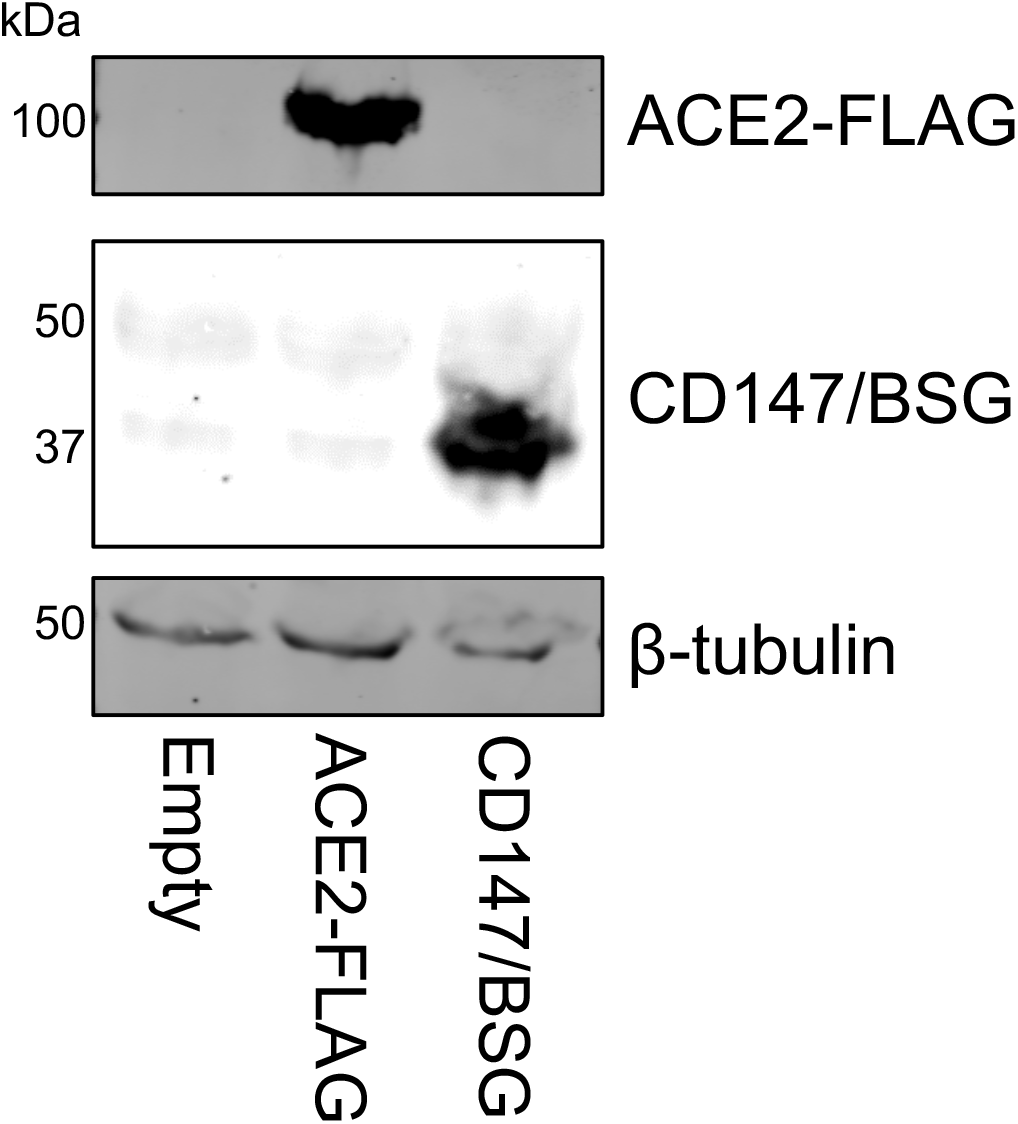
ACE2 and BSG protein expression in HEK 239Ts. Overexpression of ACE2-FLAG and CD147/BSG in HEK 293T cells was confirmed by western blot.

**Supplementary Figure 3:**
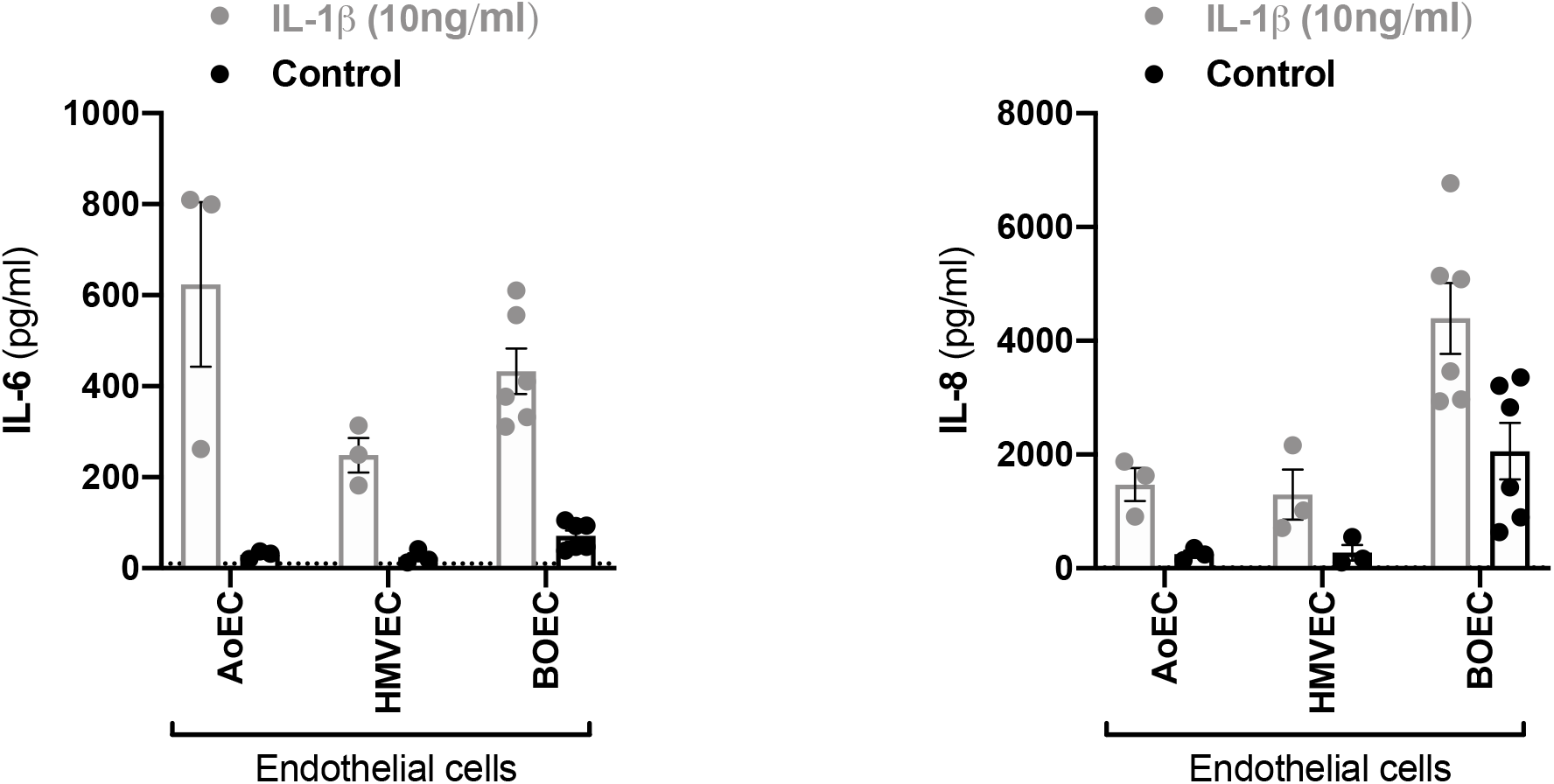
Effect of IL-1β on release of IL-6 and IL-8 from endothelial cells. Human endothelial cells (aortic; AoEC, lung microvascular; HMVEC, and blood outgrowth; BOEC) were treated for 3 hours with IL-1β (10ng/ml) before media was replaced with fresh media (EGM-2, 2% FBS) for 24 hours. Media was collected and IL-6 and IL-8 levels measured using ELISA. Data are shown as individual values and mean +/− from n=3 wells using cells from 3 separate donors for AoEC and HMVEC and n=6 wells from 3 separate donors for BOECs.

